# MAUDE: Inferring expression changes in sorting-based CRISPR screens

**DOI:** 10.1101/819649

**Authors:** Carl G. de Boer, John P. Ray, Nir Hacohen, Aviv Regev

## Abstract

Pooled CRISPR screens allow high-throughput interrogation of genetic elements that alter expression of a reporter gene readout. New computational methods are needed to model these data. We created MAUDE (Mean Alterations Using Discrete Expression) for quantifying the impact of guide RNAs on a target gene’s expression in a pooled, sorting-based expression screen. MAUDE quantifies guide-level effects by modeling the distribution of cells across sorting expression bins. It then combines guides to estimate the statistical significance and effect size of targeted genetic elements. We show that MAUDE significantly improves over previous approaches and provide experimental design guidelines to best leverage MAUDE.

## Background

Pooled CRISPR/Cas9 screens with a readout of reporter gene’s expression have improved our understanding of how both *cis* and *trans* regulators control gene expression. CRISPR/Cas9 and related systems use complementarity between an RNA guide and the genomic DNA to direct Cas9 to specific loci. Engineered Cas9 proteins with different enzymatic and regulatory activities now allow researchers to mutate DNA^1^, modify the chromatin state^2^ of the targeted locus, inertly bind loci to inhibit transcription^3^, or recruit activators^4^. Using these tools, pooled screens with libraries of guides targeting genes or the non-coding genome and measuring the impact on a reporter gene’s expression can identify *trans* regulators or *cis*-regulatory elements that affect the reporter’s expression^1, 2, 5-7^. For instance, CRISPR interference (CRISPRi) and CRISPR activation (CRISPRa) have been used to respectively repress or activate elements in a targeted *cis*-regulatory region and identify enhancers of a gene’s expression^2, 4, 8, 9^. In each case, we estimate how each guide affects the targeted (or reporter) gene’s expression by sorting cells into discrete expression bins, and measuring the guide’s abundance in the bins. Such screens help dissect enhancer regulatory logic^5, 8^, and identify the genetic variation most likely to contribute to common human disease^9^.

Despite the growing number of gene expression CRISPR screening strategies, computational methods to determine which guides and elements alter gene expression remain relatively *ad hoc*. In general, detecting guides that impact expression relies on finding those with a biased distribution across the expression bins. To date, most methods fall into two categories: (**1**) using the log fold-change between the guide abundances in the high and low expression bins directly^5, 6, 9^, or (**2**) repurposing RNA-seq differential expression tools^10, 11^ to find changes in guide abundance between high and low bins^6, 12^. However, both strategies are difficult to apply if more than two sorting bins are used and do not leverage the unique character of these data.

Here, we describe MAUDE (Mean Alterations Using Discrete Expression), an analytical framework for estimating the effect of CRISPR guides (or other perturbagens) on expression, as measured by sorting into discrete expression bins and sequencing. MAUDE maximizes the likelihood of the observed sequencing data over the cell sorting bins to estimate the mean expression for cells containing each guide. It then combines the resulting Z-scores from multiple guides targeting the same element into element-level effect sizes and estimates statistical significance. Elements can defined *a priori*, by annotating guides with the element they target, or identified *de novo* by using a sliding window to combine guides targeting neighboring chromosomal locations. MAUDE is highly sensitive at finding expression-altering guides and elements, is adept at estimating effect sizes, and outperforms existing approaches. Finally, we provide guidance on the design of expression-based CRISPR screens to maximize the information gleaned.

## Results

### MAUDE identifies expression-altering guides by their distribution across bins

To identify guides that alter expression, MAUDE makes four assumptions: (**1**) the perturbations do not differentially affect cell viability; (**2**) expression is distributed approximately (log) normally; (**3**) guides can alter the mean expression, but not the variance; and (**4**) sampling by sequencing of CRISPR guides integrated in the genomic DNA of sorted cells is captured by a negative binomial distribution (as described previously^10, 13, 14^). Note that when assumptions (2) or (3) are violated, such as if expression is bimodal (*e.g.* perturbations alter the fraction of expressing cells), MAUDE guide-level effect sizes correspond to the *average* effect of the perturbation (see below). Additionally, we require that the bin sizes be known (as a fraction of the total expression distribution of sorted cells), that the unsorted input library be sequenced to quantify overall library composition, and that the screen includes non-targeting control guides that should not affect expression.

MAUDE estimates the most likely mean expression level effect of a given guide 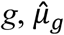, by maximizing the likelihood of the observed reads per bin under these assumptions (**Fig. 1**). To do so, it estimates the number of reads of guide *g* expected to be in each expression bin *b*, for a given mean expression level *µ*_*g*_. It estimates the optimal 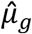 as the one that maximizes the log likelihood of the observed number of reads given the expected number of reads in each bin (**Fig. 1A** - left).

**Figure 1:**
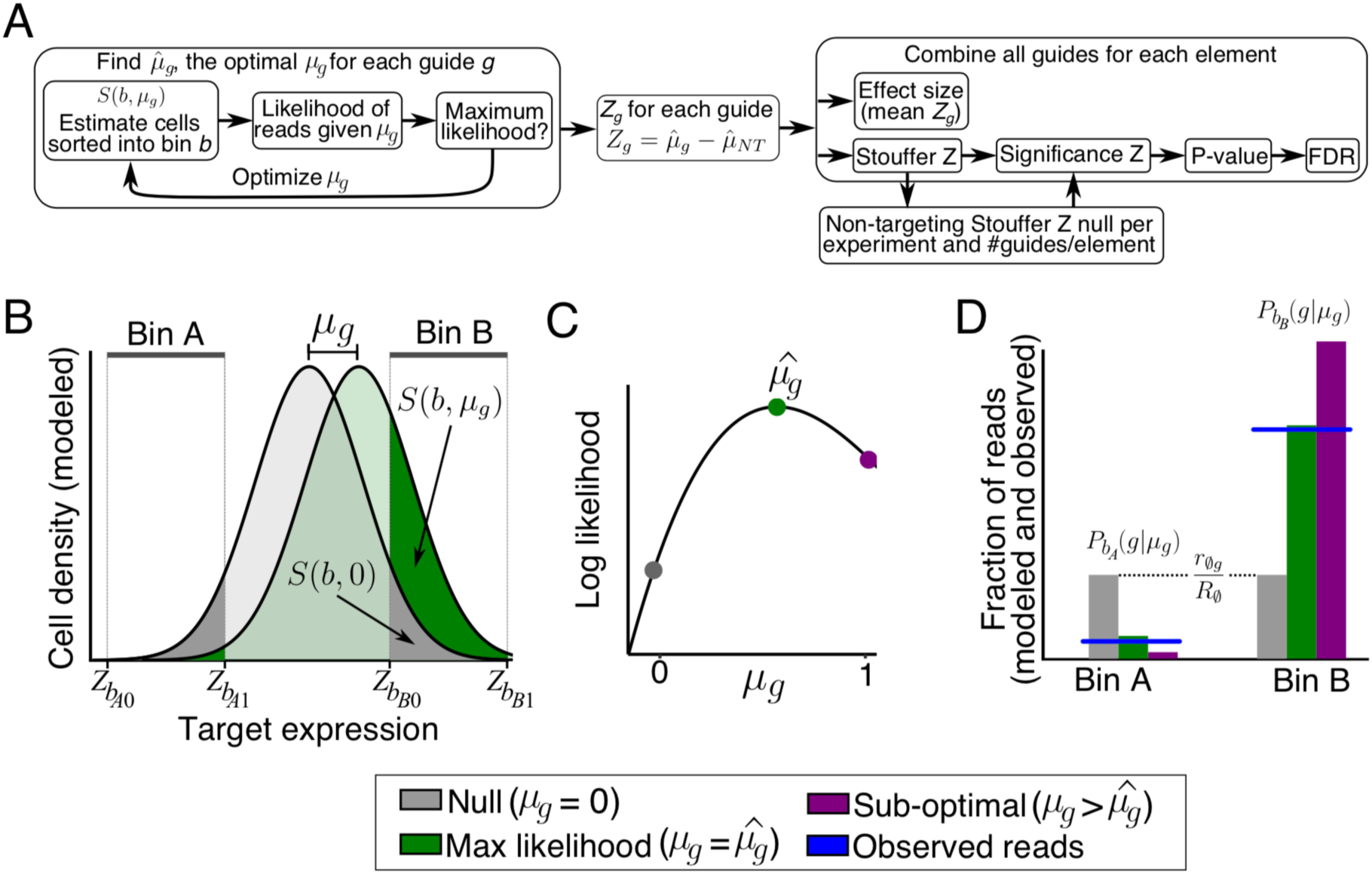
MAUDE approach to scoring expression-based screens. (**A**) Method overview. (**B-D**) MAUDE approach to estimating the optimal mean expression level per guide. (**B**) Estimation of cell density distribution for each guide. Cell density distribution (*y* axis) of target gene expression (*x* axis) for each guide is modeled as a normal distribution, with mean *µ*_*g*_. *µ*_*g*_ is optimized by calculating, for each bin (Bin A and B, top), the fraction of cells expected to be in the bin under the null model (no overall change in expression; *S*(*b*, 0)) and the fraction of cells expected in the bin given the current value of *µ*_*g*_ (*S*(*b, μ*_*g*_)). (**C**) Estimation of optimal *µ*_*g*_. We find the optimal 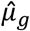 by calculating the log likelihood (*y* axis) of the observed bin read abundances given each value of *µ*_*g*_ (*x* axis). (**D**) Expected number of guide reads per bin. The fraction of reads (*y* axis) of guide *g* (*P*_*g*_(*g*|*μ*_*g*_) expected in each bin *b* (*x* axis), for different values of *µ*_*g*_ (colors).

To achieve this, MAUDE proceeds in several steps. First, it uses the unsorted (input) library to estimate the library composition, defined as *r*_ø*g*_/*R*_ø_, the fraction of reads from guide *g* (*r*_ø*g*_) out of the total reads (*R*_ø_). Next, using the overall fraction of *cells* observed in each expression bin (recorded during cell sorting), it quantile normalizes the sorting expression space to a standard normal reference (µ=0 and σ=1). For each bin, it calculates the Z-scores in standard normal space (*Z*_*b0*_ and *Z*_*b1*_ for lower and upper limit, respectively) that capture the same quantile as the bin captured in the original (FACS) data. Since most guides in such screens do not alter expression, the vast majority of the overall distribution represents the expression σ associated with no effect (*e.g.* wild type). Thus, for modeling simplicity, we can assume that each guide has the same σ – that of the overall distribution. It then defines *S*(*b, µ*_*g*_) as the fraction of cells containing guide *g* expected by the model to have been sorted into bin *b* (with the bin defined by *Z*_*b0*_ and *Z*_*b1*_, respectively). *S*(*b, µ*_*g*_) is calculated using the normal cumulative distribution function, *CDF*_*norm*_(), yielding the probability that a cell sampled from a normal distribution with mean *µ*_*g*_ will have an expression (*x*) less than the sorting upper bin bound *Z*_*b1*_, but not smaller than the lower bin bound *Z*_*b0*_:

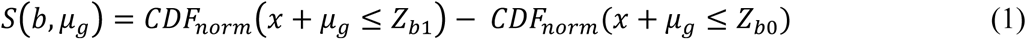

For a given *µ*_*g*_, MAUDE calculates the expected expression distribution (**Fig. 1B**). The expected fraction of cells in bin *b* that contain guide *g* is:

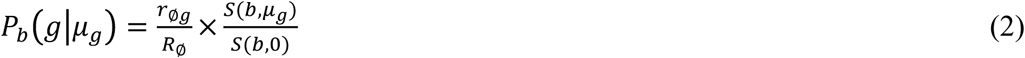

Intuitively, if guide *g* does not affect expression, *µ*_*g*_ is 0, and so *P*_*g*_(*g*|*μ*_*g*_) is just the fraction of the overall library occupied by the guide (*r*_ø*g*_/*R*_ø_); if the value of *µ*_*g*_ results in a higher fraction of cells containing guide *g* in bin *b*, the guide is enriched in that bin (*P*_*g*_(*g*|*μ*_*g*_) > *r*_ø*g*_/*R*_ø_; **Fig. 1B** - right); depletion results when fewer cells end up in the bin (*P*_*g*_ *g μ*_*g*_ < *r*_ø*g*_/*R*_ø_; **Fig. 1B** - left).

Finally, MAUDE learns the optimal *µ*_*g*_ given the observed guide reads, by maximizing the (log) likelihood of observing the reads from the guide *g* in each bin *b* (*r*_*gb*_), given the overall number of reads observed in that bin (*R*_*b*_; **Fig. 1C,D**). To reduce the impact of cases where a guide cannot be quantified reliably due to low coverage, MAUDE also includes a prior favoring no change in expression via a pseudocount added uniformly to all expression bins (**Methods**). We assume reads are sampled using a negative binomial distribution (with probability density function *PDF*_*NB*_())^10, 13, 14^. Thus, MAUDE maximizes the following equation separately for each guide *g* to learn its optimal 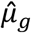:

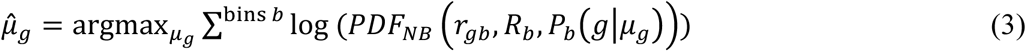

Once MAUDE learned 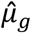 for each guide, it subtracts from each the average 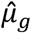 of the non-targeting guides 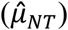 to convert each guide’s effect on expression to a Z-score (*Z*_*g*_; **Fig. 1A** – middle):

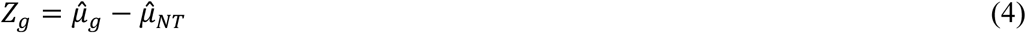

This step is necessary because targeting guides, whose true mean is sometimes not 0, are also included in the overall distribution and may have shifted the overall mean. Thus, *Z*_*g*_ represents the number of standard deviations (SDs) by which guide *g* has altered gene expression compared to the non-targeting guides.

### MAUDE calculates element level effect size and significance by aggregating signal across guides

With a Z-score representing the expression change for every guide, MAUDE calculates element-level effect size and statistical significance (**Fig. 1A** – right), using either (**1**) known element annotations or (**2**) a sliding window across the locus of interest to define elements. Examples of potential annotated elements are putative enhancers or protein coding genes. Sliding windows require that the region be tiled with guides, with the final resolution and sensitivity proportional to the tiling density, but require no prior knowledge about the region. We combine all guides within a pre-specified window size (*e.g.* 500 bp), requiring a minimum number of guides per window (*e.g.* 5), and testing all possible windows with distinct guide sets. The element’s effect size, expressed as a Z-score, is the average Z-score of the guides targeting that element.

To estimate statistical significance, we combine guide-level Z-scores for all guides targeting that element (by Stouffer’s method) into a single Z-score, representing a signed significance of the regulatory change (Z>0 representing upregulation, and Z<0 representing downregulation). Because experimental noise and the number of guides per element can greatly affect the robustness of these Stouffer Z-scores, we repeatedly sample the non-targeting guides for each experiment and every possible number of guides/element, to create null distributions. We then scale the Stouffer Z-scores by the standard deviation of the corresponding null (*i.e.* the null for that experiment with the same number of guides/element). Finally, the resulting significance Z-scores are converted into P-values using the normal cumulative distribution function (µ=0, σ=1), for the upper tail (upregulation), lower tail (downregulation), or the minimum of the two (either), with corresponding Benjamini-Hochberg FDR correction.

### MAUDE correctly identifies differential elements in simulated data

To test MAUDE’s performance and underlying assumptions, we next simulated two experiments, each with 200 targeted elements and 5 guides per element. We compared two ways in which expression might be altered by targeting of an element: (**1**) a shift from one unimodal distribution of expression levels to another, reflecting an impact on expression affecting each perturbed cell (*µ*_*g*_ ranging from 0.01 to 1 SDs, in 0.01 increments; “mean-altering”; **Fig. 2A**); or (**2**) having two fixed distributions of expression levels, as would be the case in two sub-populations, with the perturbation causing a shift in the relative proportion of cells in each expression mode (*µ*_*g*_ = 1; and the fraction of affected cells *r*_*g*_ ranges from 1% to 100%, in 1% increments; “proportion-altering”; **Fig. 2B**). In either case, 100 “effective” elements have an impact, and the other 100 “ineffective” elements have no effect on expression. We generated these synthetic data, simulating sorting cells by expression into six different 10% bins on the extremes of expression (**Methods**).

**Figure 2:**
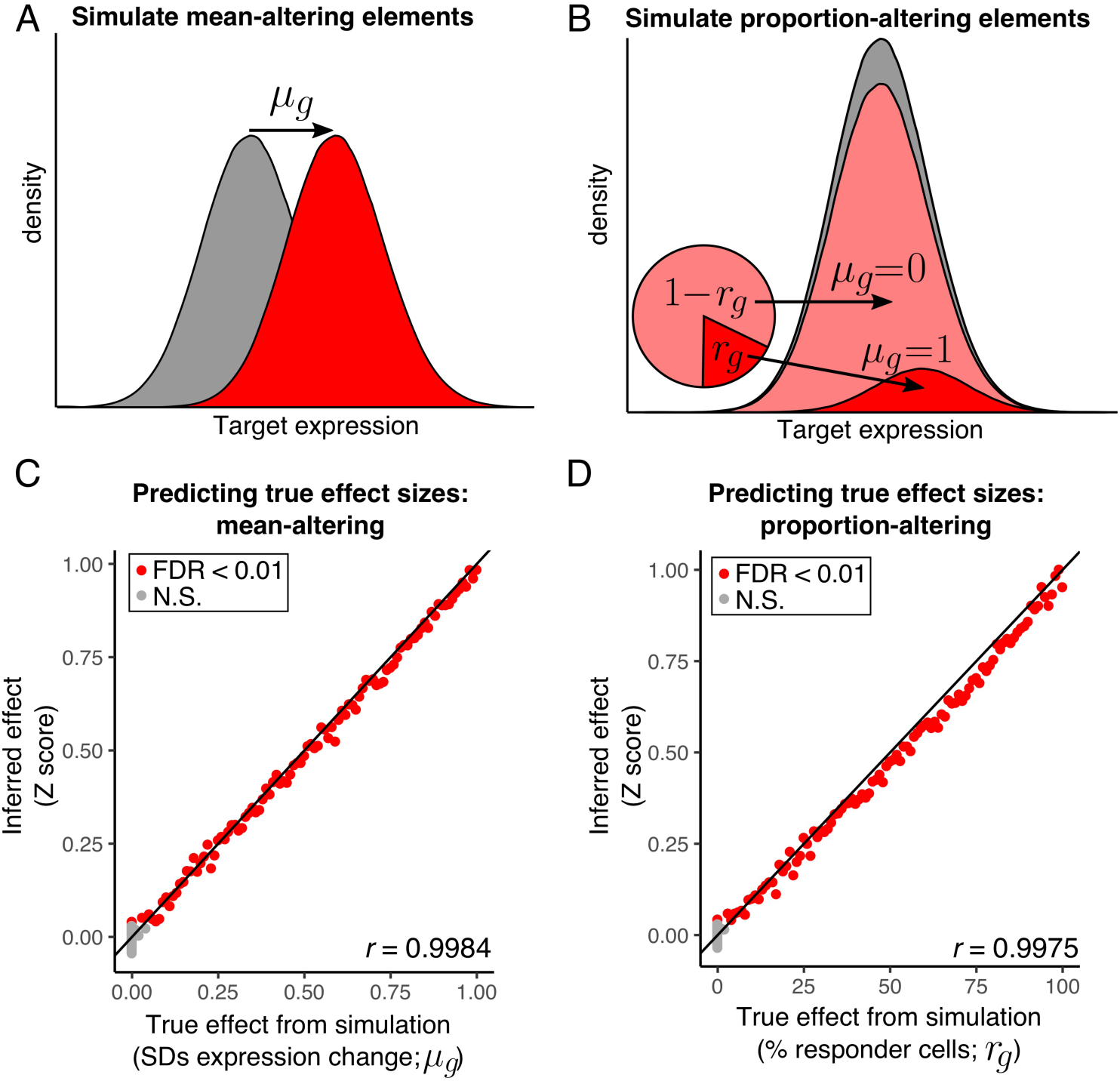
MAUDE correctly identifies functional elements and effect sizes in simulated data. (**A**) Simulation for mean-altering elements. A mean altering element is modeled as changing the mean expression *µ*_*g*_ (between 0.01 and 1) of cells with the effective element (red curve) compared to cells with ineffective elements or non-targeting guides (gray curve). (**B**) Simulation of proportion-altering elements. A proportion-altering element is modeled as resulting in a different proportion of cells with altered expression (*r*_*g*_), from 1% to 100%. On average, *r*_*g*_% of cells would have expression increased by *µ*_*g*_= 1 (dark red curve), and (1-*r*_*g*_)% of cells would have no change in expression (*µ*_*g*_ = 0; light red curve) relative to ineffective elements and non-targeting guides (gray curve). (**C, D**) Effect sizes are correctly estimated. The true effect used to generate the simulated data (*x* axis) *versus* the inferred effect size (*y* axis) for (**C**) mean-altering elements (where the true effect is *µ*_*g*_), and (**D**) proportion-altering elements (where the true effect is *r*_*g*_). Elements meeting a 1% FDR are shown in red and others in gray. Pearson’s *r* is shown in lower left (considering effective elements only). Black line: *y*=*x*.

MAUDE analysis was adept at identifying effective elements, and accurately estimated effect sizes and fractions of responding cells. In both simulations, it detected as significant (1% Benjamini-Hochberg FDR) 97 of the 100 effective elements, with effect sizes as low as 0.03 SDs (mean-altering; **Fig. 2C**) or 3% of cells responding (proportion-altering; **Fig. 2D**), indicating that MAUDE is sensitive to even small changes. As expected, each simulation included 1-2 false positives (2/99 and 1/98 for mean-altering and proportion-altering simulations, respectively; estimated effect size of ∼0.04 SDs for both), consistent with our 1% FDR, indicating that our P-values are well-calibrated. The correlation between inferred and actual effect size was very high (Pearson’s *r* = ∼0.998 for both simulations; only considering the 100 simulation-defined effective elements). In the proportion-altering simulation, the estimated effect size corresponds to the average affect size (*e.g.* 10% of cells shifting by 1 SD results in an average effect size of 0.1). Overall, MAUDE performs extremely well on synthetic data and can identify perturbations that shift the mean expression or the fraction of expressing cells with high sensitivity and specificity.

### MAUDE application to a CD69 CRISPRa screen increases accuracy and highlights new regulatory elements

We next evaluated MAUDE’s performance on actual data from a CRISPRa tiling screen to identify regions that could regulate CD69 in Jurkat cells^9^, which was conducted in duplicate. We compared MAUDE to the original analysis (“original”), where guide effects were estimated as the log ratio of normalized counts within the “high” enriched bin and “baseline” bin^9^, and with edgeR^10^ and DESeq2^11^, looking for differential guides between the “high” and “baseline” bins following previous studies^6, 12^.

We assessed the sensitivity and accuracy of each approach by three criteria, where possible: (**1**) higher similarity between effect sizes of adjacent guides, which are expected to more often target the same regulatory element, compared to pairs of randomly selected guides; (**2**) ability to distinguish promoter-targeting guides from other targeting guides, because CD69 promoter-targeting guides are a positive control expected to greatly alter CD69 expression; and (**3**) similarity between the effect sizes estimated for each replicate, which should be high if a method is successful.

MAUDE performed well on all three measures, out-performing the other approaches (**Fig. 3A-C**). MAUDE most easily distinguished adjacent from randomly paired guides (by AUROC; **Fig. 3A**), with adjacent guides having more similar effect sizes than randomly paired guides, as expected; MAUDE most easily distinguished promoter-targeting guides from other genome-targeting guides (by AUROC; **Fig. 3B**); and the estimated guide-level effects were much more correlated between the two replicates for MAUDE than for the original approach (Pearson’s *r* = 0.66 *vs*. 0.25, respectively; edgeR and DESeq2 do not return replicate-level statistics; **Fig. 3C**).

**Figure 3:**
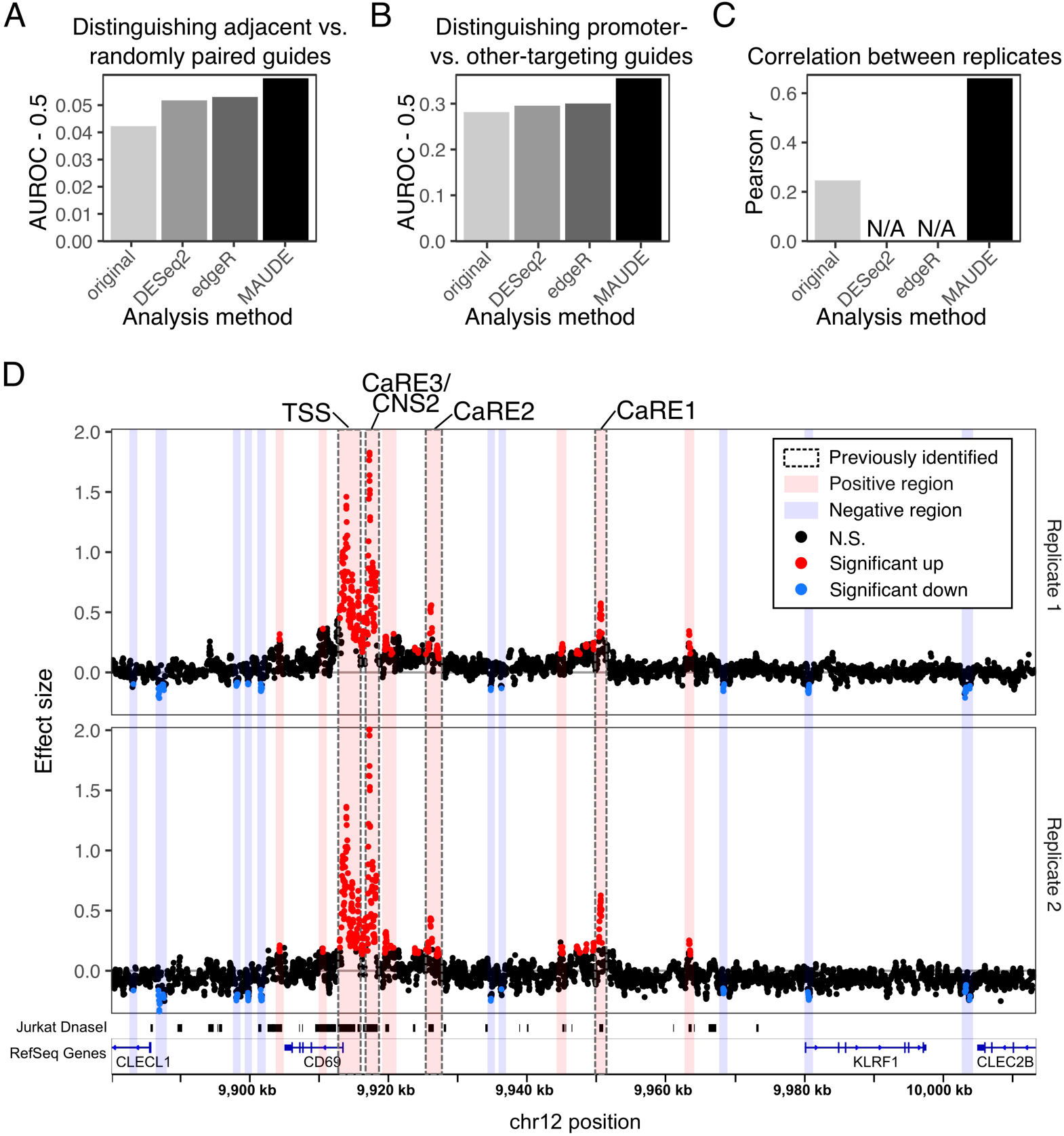
MAUDE outperforms other methods and yields insight into CD69 regulation. (**A-C**) MAUDE outperforms other methods by three criteria. Comparison of the original analysis of CD69 CRISPRa tiling screen^9^ (“original”), edgeR, DESeq2, and MAUDE (*x* axes), by three evaluation criteria (*y* axes, higher values correspond to better performance): (**A**) relative similarity in effect sizes of adjacent guides (compared to randomly selected pairs of guides, by AUROC); (**B**) ability to distinguish promoter-targeting guides from other targeting guides; and (**C**) Pearson correlation between the effect size estimates of both replicates. Correlation between replicates was not possible for edgeR or DESeq2 as they combine replicates to estimate effect sizes and statistics. (**D**) Reanalysis of CD69 CRISPRa screen data with MAUDE’s sliding window approach highlights additional plausible regulatory elements. Effect size (*y* axes) *vs* genomic position (*x* axis) for two replicates (top and bottom), with elements (points) colored by those that significantly increased (red) or decreased (blue) expression of CD69 and those that did not (black). Bottom: Gene annotations (blue) and Jurkat DNaseI hypersensitivity (black). Vertical bars: Clusters of significant elements identified by MAUDE: regions that increase CD69 expression (red), and regions that decrease CD69 expression (blue), with previously identified regions indicated (gray border).

We next asked if MAUDE yielded any new insights into CD69 regulation, which were not highlighted in the original analysis. We generated element-level statistics using a 200 bp sliding window across the locus, combining the two experimental replicates by requiring elements to have FDR < 0.01 in both replicates and consistent expression changes. The minimum element effect size that was reproducibly found by MAUDE was a change of 0.12 SDs.

MAUDE re-discovered the four regions previously called as being CRISPRa-sensitive (**Fig. 3D**, gray bars), and called 15 additional regions as responsive to CRISPRa. Of these, 10 appeared to cause a *down*-regulation of CD69 when activated by CRISPRa (**Fig. 3D**, red bars). Although none of these are in open chromatin regions in Jurkat cells, two are adjacent to the promoters of other nearby genes (CLECL1 and CLEC2B), suggesting these may act by competition with the CD69 promoter. Similar opposing effects on expression have been previously reported when targeting the promoters of neighboring genes with CRISPRi^8, 12^. The remaining five CRISPRa-sensitive regions we identified caused upregulation of CD69; all five of these regions overlapped with Jurkat open chromatin regions (**Fig. 3D**, red bars), as did the CRISPRa-sensitive regions originally identified^9^. Overall, 85% of CD69-upregulating elements identified by MAUDE were within Jurkat open chromatin and were closer to open chromatin than expected by chance (rank sum P<10^−15^; distances to open chromatin *vs*. distances to randomly-placed open chromatin; **Methods**).

### MAUDE helps design bin number and width to enhance screen sensitivity

While an experimenter can set the sizes of the expression bins, finding an optimal bin configuration experimentally is laborious and costly, and typically not pursued in practice. We reasoned that our simulation framework can readily test different expression bin configurations to help choose the best one under our assumptions.

To this end, we accounted for several experimental considerations. First, cell sorters can have between two to six bins available, and the optimal bin size may differ when different numbers of bins are used. Second, the tails of the distribution are the most informative because they show the greatest ratios of effective versus ineffective guides, and this ratio increases as bins get more extreme. However, the more extreme the bins, the fewer cells will be captured, resulting in higher sampling noise. Finally, it is desirable to have each bin capture approximately the same number of cells to facilitate uniform treatment of samples (*e.g.* genomic DNA isolation and library preparation).

With these considerations in mind, we tested by simulation how bin size and number altered the ability to identify differentially active elements (**Fig. 4A**). We altered our simulation framework to include more elements (1,000 ineffective and 1,000 effective), and, to make the recovery of effective elements more challenging, focused on smaller effect sizes (100 each with effect sizes from 0.01 to 0.1, at 0.01 increments). Here, the smallest effect sizes are the most difficult to recover and only the most sensitive approaches will recover them. Our bin configurations included both uniform percentile bins and more complex bin configurations, always placing bins symmetrically at both tails of the distribution, and, within each tail, bins are contiguous (**Fig. 4A**, right). Uniform binning schedules were also tested with two and four bins, always retaining the most extreme bins. Otherwise, six bins were used. We evaluated bin configurations for their accuracy in predicting effect sizes (Pearson’s *r*) and their sensitivity in recovering functional elements (true positives).

**Figure 4:**
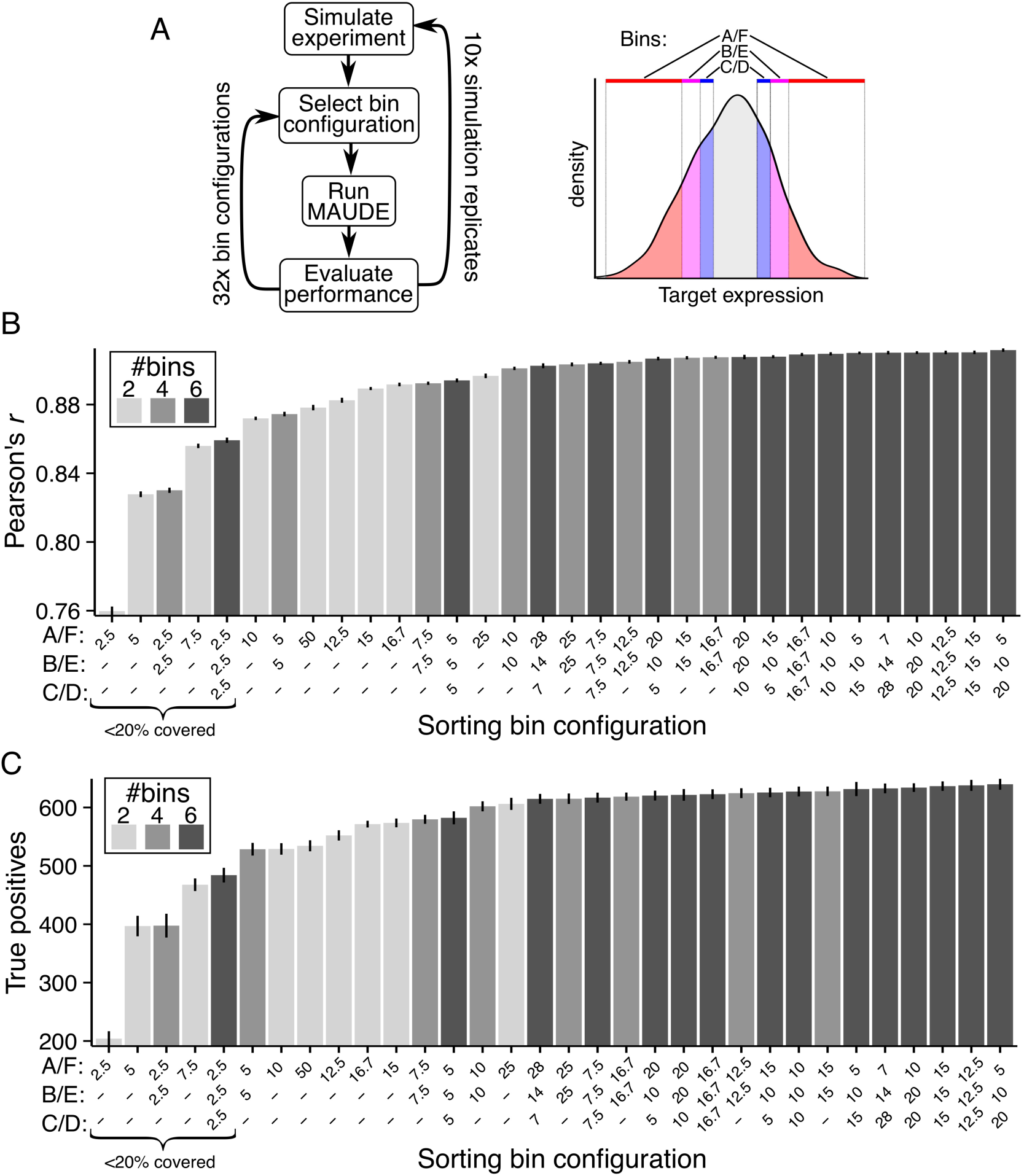
MAUDE simulations propose optimal cell-sorting bin configurations. (**A**) Evaluation of expression bin configurations by simulation. Left: An experiment is simulated (10 replicates, top), followed by selection of each bin configuration, MAUDE analysis and evaluation of performance for true positive recovery and effect size prediction (by Pearson’s *r*). Right: Each bin configuration had 2-6 bins that are contiguous on each end and symmetric about the center, progressing gradually with inclusion from outer to innermost: A and F: outermost and always included, B and E (when present): contiguous with A and F (respectively) and more central; and C and D (when present): contiguous with B and E (respectively) and innermost. (**B, C**) Identification of optimal bin configurations. Performance (*y* axis) of each bin configuration (*x* axis) for (**B**) Pearson’s *r* between the true effect sizes (as defined in the simulation) and the predicted effect sizes (by MAUDE), and (**C**) the number of true positives (elements defined as having an effect in the simulation that are predicted to have an effect by the analysis). *x* axis labels: Bin sizes expressed as the percent of the distribution covered by each of the paired corresponding bins, sorted in order of increasing performance. Error bars: standard error of the mean (SEM) for the 10 simulated experiments. Gray shading: number of bins in the configuration.

The best bin configurations include more bins and cover of most of the distribution. When using non-uniform bins, greater resolution (*i.e.* smaller bin sizes) at the tails of the distribution (*i.e.* A/F scheme, **Fig. 4A**) resulted in both greater sensitivity and accuracy than greater resolution at the inner bins (**Fig. 4B,C**). Having more sorting bins was always better, but particularly when bins are small, and going from two to four bins was a much more significant increase than from four to six bins (**Fig. 4B,C**). Overall, performance was relatively poor when bins covered less than 20% of the overall distribution (**Fig. 4B,C** – indicated at left). However, performance is also poor if bins are too large, presumably because resolution is reduced. For example, having two 50% bins was worse than two 25% bins, and four 25% bins was worse than four 12.5% bins (**Fig. 4B,C**). The best configuration tested by both measures had A and F cover 5%, B and E cover 10%, and C and D cover 20%, but uniform bins covering 12.5 - 15% had similar performance (**Fig. 4B,C**) and yield uniform cell numbers per sample, and so are more practical. Overall, this simulation indicates that the greatest sensitivity and accuracy is achieved by having more bins, good performance can be achieved with uniform bins, and each of the bins should cover about ∼15% of the distribution (∼25% if using only two bins).

## Discussion

MAUDE is a principled framework for the analysis and design of pooled CRISPR screens with an expression readout. By estimating the effects of guides and elements from the distribution of guides across bins, we show that MAUDE is both sensitive and specific on simulated data, and more accurate on real data, allowing the recovery of additional regulatory elements. Notably, although our sliding window approach can be used for identifying regulatory regions in a tiling experiment, MAUDE can also be used in combination with other approaches^15^. MAUDE’s simulation framework can help experimental design to maximize a screen’s success.

Although we focus our testing on CRISPR screens aiming to identify regulatory elements with a guide RNA perturbagen, which are the current common examples, we expect any experiment with a binned expression readout and sequencing to quantify perturbations could use MAUDE analysis. This could include perturbations of genes ^6^, other types of perturbagens (*e.g.* RNAi), direct readouts of mutation (*e.g.* via base editors), or even reporter assays^16^.

## Conclusions

MAUDE is a highly sensitive and accurate approach for identifying functional elements in a binned expression screen. MAUDE estimates the effect size on expression of individual perturbagens (*e.g.* CRISPR guide RNAs), and combines perturbagens to estimate the effects of genetic elements. MAUDE performs well in simulations, and outperforms other methods on experimental data by all three independent evaluation criteria. Finally, by simulation, we identify which cell-sorting bin configurations are optimal.

As research focused on the function on regulatory variants and regions, which are disproportionately contributing to the genetic basis of common disease^17^, we expect that pooled genetic screens with expression readouts will become much more common. Moreover, many such variants and elements are expected to have smaller effect sizes, emphasizing the need for sensitive detection and accurate effect size estimates. We anticipate that MAUDE – which we implemented as an R package available on GitHub (https://github.com/Carldeboer/MAUDE) – will become an important tool in the effort to map from variants to function.

## Methods

### Implementation and usage

MAUDE is implemented in R. Tutorials are provided on the MAUDE website (https://github.com/Carldeboer/MAUDE). Users provide a data.frame, with columns containing the bin counts (one column per bin, plus one for the unsorted cells), as well as columns annotating the data included in each row, including: guide ID, experimental identifiers (*e.g.* replicate, condition, etc.), whether or not the guide is a non-targeting guide, and any other guide-associated data (*e.g.* genomic locus). Users also provide a data.frame containing the bin sizes, with one row per bin per experiment, and columns corresponding to the Z-score bounds of each bin (‘binStartZ’ and ‘binEndZ’) and the corresponding bin cumulative distribution function percentiles (‘binStartQ’ and ‘binEndQ’). Using MAUDE’s ‘findGuideHitsAllScreens’ function, and providing the experimental design data.frame, read count data.frame, and bin bound data.frame, MAUDE will calculate the optimal mean expression for each guide, separately for each experiment, returning the mean guide expression 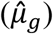, Z score (*Z*_*g*_), and log likelihood ratio for each guide/experiment pair. By default, for each guide *<u>g</u>* and bin *b*, 10 pseudocounts are added to the read count (*r*_*g,b*_) for every million reads of coverage in that bin (*R*_*b*_) to reduce the noise resulting from poor coverage. R’s ‘optimize’ function is used for finding the guide mean expression with the highest log likelihood. The fraction of cells expected to be in a bin given the current *μ*_*g*_ is calculated with the ‘pnorm’ function, scaled by the overall abundance in the library of that guide and the fraction of cells sorted into the bin, as in Equation (2). The log likelihood for each guide/bin is calculated as using the ‘dnbinom’ function, with x = number of reads for this guide in this bin, size = number of reads total for this bin, prob = 1 - the fraction of this bin expected to be occupied by this guide, as in Equation (3). The ‘findGuideHitsAllScreens’ will return the guide-level statistics as a data.frame.

Once guide-level statistics have been calculated for each experiment, they can then be combined to obtain element-level statistics. Using MAUDE’s ‘getTilingElementwiseStats’ function, one can identify elements *de novo* by tiling across the region, providing the experimental design data.frame and the guide-level statistics data.frame, including columns denoting the genomic coordinates of the guide (chromosome, position). By default, all guides within each 500 bp window are combined in this sliding window approach, requiring a minimum of five guides per window (parameters which can and should be customized, dependent on the density of tiled guides), and testing all possible windows with unique guide sets. Alternatively, MAUDE provides the ‘getElementwiseStats’ function to calculate element-level statistics given element annotations as an additional column in the guide-level statistics data.frame.

In either case, element-level statistics are calculated by combining all Z-scores for guides in the element. Three Z-scores are calculated for each element: (1) an effect size – the mean of the guides’ Z-scores; (2) the Stouffer Z-score – the guides’ Z-scores combined with Stouffer’s method (‘stoufferZ’); and (3) a significance Z-score – the Stouffer Z-score scaled using the appropriate non-targeting null model (‘significanceZ’). To create robust null models used to scale the statistical Z scores, we sample each non-targeting guide up to 10 times each. For each experiment and number of guides per element, we calculate the standard deviation of the null Stouffer Z scores. By dividing by this standard deviation, we ensure that the null now has a standard deviation of 1, and so we can treat them as a true Z scores, calculating P-values with the ‘pnorm’ function. P-values can be calculated with one-tail (up-/down-regulation) or two tails (either). FDRs are then calculated per-experiment using the Benjamini-Hochberg procedure.

### Synthetic data generation

For each simulation, we included 1,000 targeting guides (5 per element for 200 elements) and 1,000 non-targeting guides designed to have no effect. The underlying abundance of each guide (*A*_*g*_) followed a Poisson distribution with mean 1,000 (representing library construction noise; *A*_*g*_ ∼ *Pois*(1000)). The number of cells sorted for each guide (*S*_*g*_) followed another Poisson distribution with a mean corresponding to the abundance of that guide within the library (*S*_*g*_ ∼ *Pois*(*A*_*g*_)). For the mean-altering simulation (**Fig. 2A,C**), expression levels of cells (*E*_*gc*_) were simulated by sampling from a normal distribution with a mean corresponding to the assigned mean expression of that guide (*E*_*gc*_ ∼ *Norm*(*µ*_*g*_)) for each sorted cell *c*. For the proportion-altering simulation (**Fig. 2B,D**), cells were partitioned into responders and non-responders with a probability proportional to the assigned fraction of cells whose expression is changed by the perturbation (*r*_*g*_), and the expression followed a normal distribution with a mean 0 for non-responder cells and 1 for responder cells. The number of cells “sorted” into each bin (*S*_*bg*_) using the bounds of each bin is:

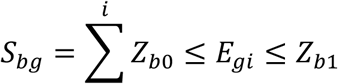

Reads were simulated with a negative binomial distribution with the number of reads for each bin equal to ten times the number of cells sorted into that bin and the probability of selecting a guide equal to its fraction of cells within that bin (**Fig. 2C**):

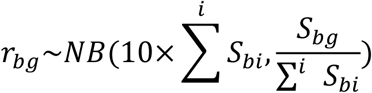

### MAUDE, logFC, edgeR, and DESeq2 application to CD69 data

Raw count data for the CD69 CRISPRa screen was downloaded from PubMed Central (Simeonov et al. Supplementary Table 1)^9^. The logFC values of the original analysis were used as-is (“original”). EdgeR (v 3.16.1) analysis was done with default parameters by comparing high to baseline bins, for each of the two replicates, together, normalizing the data (calcNormFactors) and performing a likelihood ratio test (estimateDisp, glmFit, glmLRT), and the significance p-values and estimated log fold change retrieved (topTags). DESeq2 (v 1.14.1) analysis was done with default parameters by creating a DESeq2 experiment using the CD69 count data, comparing the high and baseline bins, using both replicates (DESeqDataSetFromMatrix), and then comparing the expression between these bins (using the “DESeq” function). MAUDE analysis was performed with default parameters on the count data for each bin, including the unsorted bin. We estimated the fraction of cells in each bin for use in MAUDE analysis by reconstructing the expression distribution in Simeonov et al. Extended Data Fig. 1a^9^ using the “digitize” R package^18^, using the same data for both replicates. Using the actual bin proportions for each replicate would likely increase MAUDE’s performance.

For each evaluation criterion, we used effect sizes or significance values, as appropriate. Effect sizes were: the mean of the two replicates’ guide Z-scores (MAUDE); the estimated log fold change (edgeR and DESeq2); or the mean logFC for the two replicates as originally reported (“original”). Significance values were: combined replicates’ Z scores by Stouffer’s method (MAUDE); estimated p-values (edgeR and DESeq2); and the mean of the two replicates’ logFCs (“original”; no significance estimates were given in Simeonov et al.).

For testing whether adjacent guides have similar effect sizes, we sorted the guides by the genomic coordinates of the target site, and, for each pair of adjacent guides, calculated the absolute difference in effect sizes. To create the random-pairing distribution, this was repeated, including each guide 10 times (to improve our “random” estimate), and randomizing the order of the guides. The distributions in absolute effect size differences between adjacent and randomly paired guides were compared with the Area Under the ROC curve statistic (AUROC), and how much it differed from that expected by chance (0.5; *i.e.* AUROC-0.5). Here, the AUROC is proportional to the fraction of randomly paired guides whose absolute effect size differences are greater than those of adjacent guides.

The AUROC statistic was also used to compare the guide significance distributions between CD69 promoter-targeting guides and other genomic targeting guides, where the AUROC is proportional to the fraction of promoter targeting guides that are more significant than non-promoter targeting guides. Promoter-targeting guides were defined as those in the 1 kb surrounding the CD69 TSS (between 9912997 and 9913996 on chromosome 12; hg19).

For testing the correlation between replicates, the original logFC values and MAUDE guide Z-scores were used directly.

Element-level statistics for CD69 screen were calculated with the sliding window approach, using a 200 bp window and requiring a minimum of five guides per element. To estimate the significance of the overlap between CD69-activating elements and Jurkat open chromatin (DNaseI hypersensitivity), the distance distributions between open chromatin sites and MAUDE-identified CD69-activating elements were compared for actual *vs.* randomized open chromatin sites. The randomization placed open chromatin elements in between the starts of the first and last open chromatin sites in the locus (chr12: 9885788 – 9973087), accepting only random placements with no overlap between open chromatin sites, and preserving their widths. 100 such randomizations were created, each time computing the distances between the MAUDE-identified CD69-upregulating elements and the nearest randomly-placed open chromatin element. The distance distributions for the actual open chromatin data *vs.* the randomized open chromatin data were then compared using a two-tailed Wilcoxon rank sum test.

## Abbreviations

AUROC: Area under the receiver operating characteristic curve
CaRE: CRISPRa responsive element (as identified in ^9^)
CRISPR: clustered regularly interspaced short palindromic repeats
FDR: false discovery rate
MAUDE: mean alterations using discrete expression
SD: standard deviation
TSS: transcription start site

## Availability of data and software

All data analyzed during this study are included in the supplementary information of Simeonov et al. (Supplementary Table 1) ^9^. MAUDE is available as an R package on GitHub (https://github.com/Carldeboer/MAUDE).

## Authors’ contributions

CGD conceived of and implemented the algorithms with help from JPR. CGD drafted the manuscript, with input from JPR, AR, and NH. All authors read and approved the final manuscript.

## Acknowledgements

We thank Benjamin Doughty and Charles Fulco for helpful discussions.

CGD was supported by a Fellowship from the Canadian Institutes for Health Research and the NIH (K99-HG009920-01). JPR is funded by NIH 5 F32 AI129249. Work was supported by the Klarman Cell Observatory and HHMI (AR), the Center for Cell Circuits, a Center of Excellence in Genome Science (RM1HG006193), and the NHGRI (R01HG008131-01; NH). NH is the David P. Ryan endowed chair, and thanks Sandra, Sarah and Arthur Irving for support.

AR is an equity holder and founder of Celsius Therapeutics and an SAB member of ThermoFisher Scientific, Neogene Therapeutics and Syros Pharmaceuticals. NH is a co-founder and equity holder of Neon Therapeutics. All other authors declare no competing interests.

